# TAD-like single-cell domain structures exist on both active and inactive X chromosomes and persist under epigenetic perturbations

**DOI:** 10.1101/2021.05.12.443887

**Authors:** Yubao Cheng, Miao Liu, Mengwei Hu, Siyuan Wang

**Affiliations:** Department of Genetics, Yale School of Medicine, Yale University, New Haven, CT 06510, USA; Department of Cell Biology, Yale School of Medicine, Yale University, New Haven, CT 06510, USA; Yale Combined Program in the Biological and Biomedical Sciences, Yale University, New Haven, CT 06510, USA; Molecular Cell Biology, Genetics and Development Program, Yale University, New Haven, CT 06510, USA; Biochemistry, Quantitative Biology, Biophysics and Structural Biology Program, Yale University, New Haven, CT 06510, USA; M.D.-Ph.D. Program, Yale University, New Haven, CT 06510, USA; Yale Center for RNA Science and Medicine, Yale University School of Medicine, New Haven, CT 06510, USA; Yale Liver Center, Yale University School of Medicine, New Haven, CT 06510, USA

**Keywords:** Topologically associating domain (TAD), TAD-like structure, single-cell domain, X chromosome, X inactivation, chromatin folding, chromatin compaction, image-based spatial genomics, 3D genomics, chromatin tracing, multiplexed sequential fluorescence *in situ* hybridization (FISH)

## Abstract

**Background:** Topologically associating domains (TADs) are important building blocks of three-dimensional genome architectures. The formation of TADs was shown to depend on cohesin in a loop-extrusion mechanism. Recently, advances in an image-based spatial genomics technique known as chromatin tracing led to the discovery of cohesin-independent TAD-like structures, also known as single-cell domains — highly variant self-interacting chromatin domains with boundaries that occasionally overlap with TAD boundaries but tend to differ among single cells and among single chromosome copies. Recent computational modeling studies suggest that epigenetic interactions may underlie the formation of the single-cell domains.

**Results:** Here we use chromatin tracing to visualize in female human cells the fine-scale chromatin folding of inactive and active X chromosomes, which are known to have distinct global epigenetic landscapes and distinct population-averaged TAD profiles, with inactive X chromosomes largely devoid of TADs and cohesin. We show that both inactive and active X chromosomes possess highly variant single-cell domains across the same genomic region despite the fact that only active X chromosomes show clear TAD structures at the population level. These X chromosome single-cell domains exist in distinct cell lines. Perturbations of major epigenetic components did not significantly affect the frequency or strength of the single-cell domains. Increased chromatin compaction of inactive X chromosomes occurs at a length scale above that of the single-cell domains.

**Conclusions:** In sum, this study suggests that single-cell domains are genome architecture building blocks independent of the tested major epigenetic components.

## Background

Genomic DNA is compactly folded into the cell nucleus and is spatially organized with other nuclear components in eukaryotic cells^1–8^. This folding and spatial organization has been shown to control many important genomic functions, and altered organization has been linked to diseases^1–11^. Recent studies have identified topologically associating domains (TADs, also known as contact domains) as an important organizational unit of chromatin folding^12–16^. TADs are continuous sections of the genome at a length scale of around tens to hundreds of kilobases (kb) with enhanced self-interactions, initially discovered by high-throughput versions of population-averaged chromosome conformation capture methods *(e.g.* Hi-C^3^,^17^). Imaging studies showed that, in real space, individual TADs are physically separated, though adjacent TADs may partially overlap^13,18–21^. TADs are implicated in defining the scope of promoter-enhancer interactions^22,23^. Mechanistically, at least a subset of TADs are established through a loop-extrusion mechanism, in which cohesin dynamically extrudes a chromatin loop until the cohesin motor is blocked by CCCTC-binding factor (CTCF) bound to the DNA in an effective orientation^24–27^. Here the chromatin loop forms a TAD and the CTCF binding site marks the TAD boundary. In addition, TAD boundaries at least occasionally coincide with the boundaries of epigenetic domains along the genome^12,13^.

At the single-cell or single-chromosome-copy level, TADs are highly variant structures. Single-cell sequencing studies showed that the TADs identified from population averaged Hi-C maps do not form in every cell^28,29^. Recently, an image-based spatial genomics study using chromatin tracing^30,31^, a highly multiplexed sequential DNA fluorescence *in situ* hybridization (FISH) technique, showed that TAD boundaries on human chromosome 21 are highly variant from one copy of the chromosome to another^32^. To distinguish from the population averaged TAD structures, these highly variant single-cell structures were named as TAD-like structures^32^, later also named as single-cell domains^33^. Strikingly, single-cell domains persisted in single chromosomes even after the removal of key loop-extrusion machinery and after the disappearance of TAD structures at the population level^32^. Recent computational modeling studies suggested that dynamic chromatin contacts captured by epigenetic interactions may establish the single-cell domains^34,35^. Following this hypothesis, we reason that the active and inactive X chromosomes of a female cell may show distinct features of single-cell domains, given the drastically different epigenetic states of the two chromosomes despite largely identical sequences^36–38^. Allele-specific Hi-C studies showed that the inactive X chromosome is largely devoid of TAD structures, while the active X chromosome contains TADs^39,40^. It is unclear whether single-cell domains exist on both X chromosomes, and whether the single-cell domains are affected by changes in the epigenetic states. To answer these questions, here we perform chromatin tracing on the active and inactive X chromosomes in human cell lines. Our results show that single-cell domains are present on both active and inactive X chromosomes at comparable frequencies and strengths. We made this same observation in cells of distinct biological backgrounds. Chemical perturbations of several major epigenetic components do not appear to affect the single-cell domains. In addition, by comparing the fine-scale chromatin tracing data in this work with our previous work reporting larger scale chromatin traces, we also show that the increased compaction of the inactive X chromosome in comparison to the active X chromosome occurs at a length scale above that of single-cell domains. In sum, these observations offer a single-cell view of the principles of X chromosome folding and compaction, and suggest that the mechanism underlying the establishment and/or maintenance of single-cell domains may not involve the tested major epigenetic components.

## Results

To investigate the chromatin folding of active and inactive X chromosomes (Xa and Xi) at the TAD-length scale in single cells, we used chromatin tracing^30,31^ to trace the chromatin folding organization of an 840-kb region of the X chromosome in a human female IMR-90 cell line. This selected 840-kb region spans a TAD boundary, based on previously published IMR-90 Hi-C datasets^12,16^. We partitioned this region of interest into consecutive 30-kb segments, labeled and imaged each segment following our previously established chromatin tracing protocol^30,41^. In brief, we first labeled and imaged the entire region with a library of dye-labeled primary oligonucleotide probes. Each primary probe contains a targeting sequence that hybridizes to the genomic DNA, and an overhanging readout sequence that is unique to each 30-kb segment (Fig. 1A). We then sequentially hybridized to the sample dye-labeled secondary probes with sequences complementary to the readout sequences on the primary probes, allowing each 30-kb segment to be sequentially visualized using three-dimensional (3D) epifluorescence microscopy (Fig. 1A). We performed two-color imaging to simultaneously visualize two segments at a time, and then photobleached the sample before the next round of sequential hybridization. This hybridization-imaging-bleaching procedure was repeated until all secondary probes were applied and all chromatin segments were imaged (Fig. 1A). The center positions of all segments were individually measured and linked based on their order on the genomic map to reconstruct the 3D folding conformation of the region in single copies of X chromosomes in single cells (Fig. 1A). To distinguish Xa and Xi, we performed either co-immunofluorescence targeting macroH2A.1 (mH2A1), a histone variant enriched on Xi, or co-RNA FISH targeting *Xist* long noncoding RNA (Fig. 1A).

**Figure 1.**
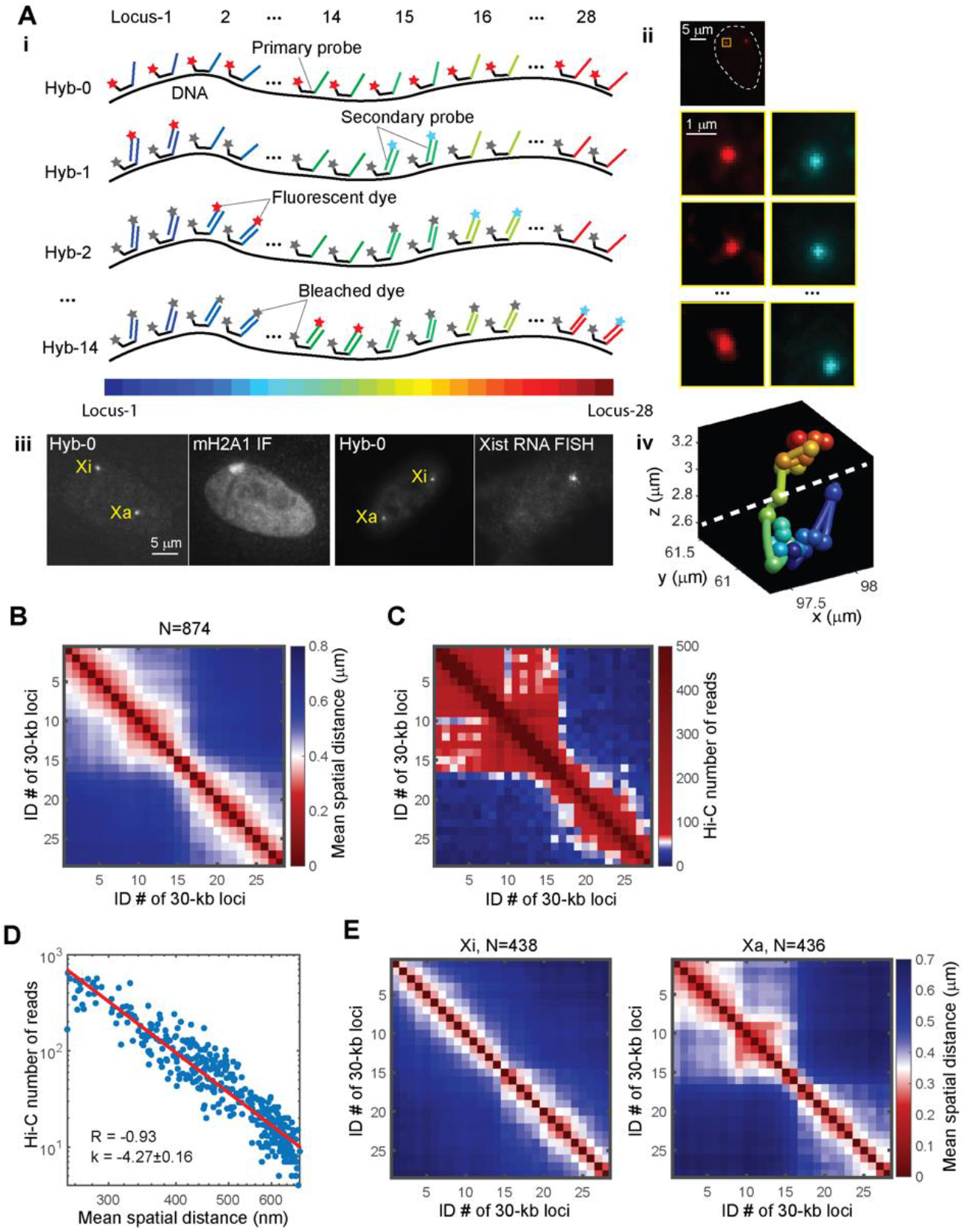
Chromatin tracing reveals the fine folding architecture of inactive and active X chromosomes. (A) Schematic illustration of the chromatin tracing approach (i), raw images of the chromatin tracing, co-immunofluorescence of mH2A1 and co-RNA FISH of *Xist* (ii and iii), and a reconstructed chromatin trace showing two domains (iv). In ii, the white dashed line in the top panel outlines a cell nucleus. The yellow boxed region containing a copy of the targeted chromatin region is shown in the panels below. The four rows of panels correspond to Hybs-0, 1, 2, and 14 as indicated on the left. (B) Mean spatial distance matrix of the traced genomic region calculated from pooled X chromosome copies including both active and inactive X’s in IMR-90 cells (N = 874). (C) Hi-C contact frequency matrix of the same genomic region as in B. (D) Comparison of mean spatial distance from chromatin tracing with contact frequency measured by Hi-C. (E) Separate mean spatial distance matrices from inactive (left, N = 438) and active (right, N = 436) X chromosomes.

To validate the chromatin traces, we calculated a mean spatial distance matrix with matrix elements representing the mean spatial distances between pairs of the targeted 30-kb segments, and compared these distances from imaging with the corresponding contact frequencies from published ensemble IMR-90 Hi-C data^16^. The mean spatial distance matrix displayed a TAD boundary at the same genomic position as in the ensemble Hi-C contact frequency matrix (Fig. 1B-C). The mean spatial distance measurements were highly correlated with the Hi-C contact frequencies, with a correlation coefficient of −0.93 (Fig. 1D). A power-law fitting showed that the Hi-C contact frequency was inversely proportional to the 4^th^ power of the mean spatial distance (Fig. 1D), similar to the power-law relationship previously observed for autosome chromatin folding^30,32,33,42^. These observations indicate that our chromatin tracing results are consistent with Hi-C results at the ensemble level, and provided a validation of the chromatin tracing data.

Next, to compare the chromatin folding conformations between Xa and Xi, we generated separate mean spatial distance matrices for Xa and Xi. The Xa matrix showed a prominent TAD boundary at the same coordinate as the whole-population TAD boundary, while the Xi matrix did not show a strong boundary (Fig. 1E). This observation is consistent with previous allele-specific Hi-C studies showing that TADs are largely attenuated on Xi^39,40^, likely due to a global loss of cohesin along Xi^39^. This consistency supports the accuracy of our identification of Xi versus Xa.

To detect potential single-cell domains in individual copies of X chromosomes, we calculated the individual spatial distance matrices for single copies of Xa and Xi, and found that individual Xa and Xi copies both displayed single-cell domains with sharp boundaries (Fig. 2A-B). The boundary positions of the single-cell domains were highly variant among individual copies of Xa and Xi (Fig. 2A-D). Quantification of boundary probabilities along the traced genomic region showed that in Xa, single-cell domain boundaries preferentially resided at the ensemble TAD boundary, whereas in Xi, the single-cell domain boundaries had relatively uniform distribution across the traced region (Fig. 2C-D), leading to the lack of strong ensemble boundary on Xi (Fig. 1E). Quantifications of the boundary strengths showed that the boundaries of single-cell domains in Xa and Xi are similarly strong (Fig. 2E-F). The single-cell domains reflect prevalent cooperativity of chromatin interaction^32^. To measure this cooperativity, we compared the conditional and unconditional contact probabilities among triplets of chromatin segments, using 200 nm as the threshold of contact: Given each combination of ordered chromatin segment A, B and C, we measured the unconditional contact probability between B and C, and compared that with the conditional contact probability between B and C given that A and B were making contact, as well as the conditional contact probability between B and C given that A and B were not making contact. The data showed that in both Xa and Xi, the conditional probability of B-C contact given A-B contact is systematically higher than the unconditional probability, and higher than the conditional probability without A-B contact (Fig. 2G). These results indicate that both Xa and Xi contain single-cell domains of similar features.

**Figure 2.**
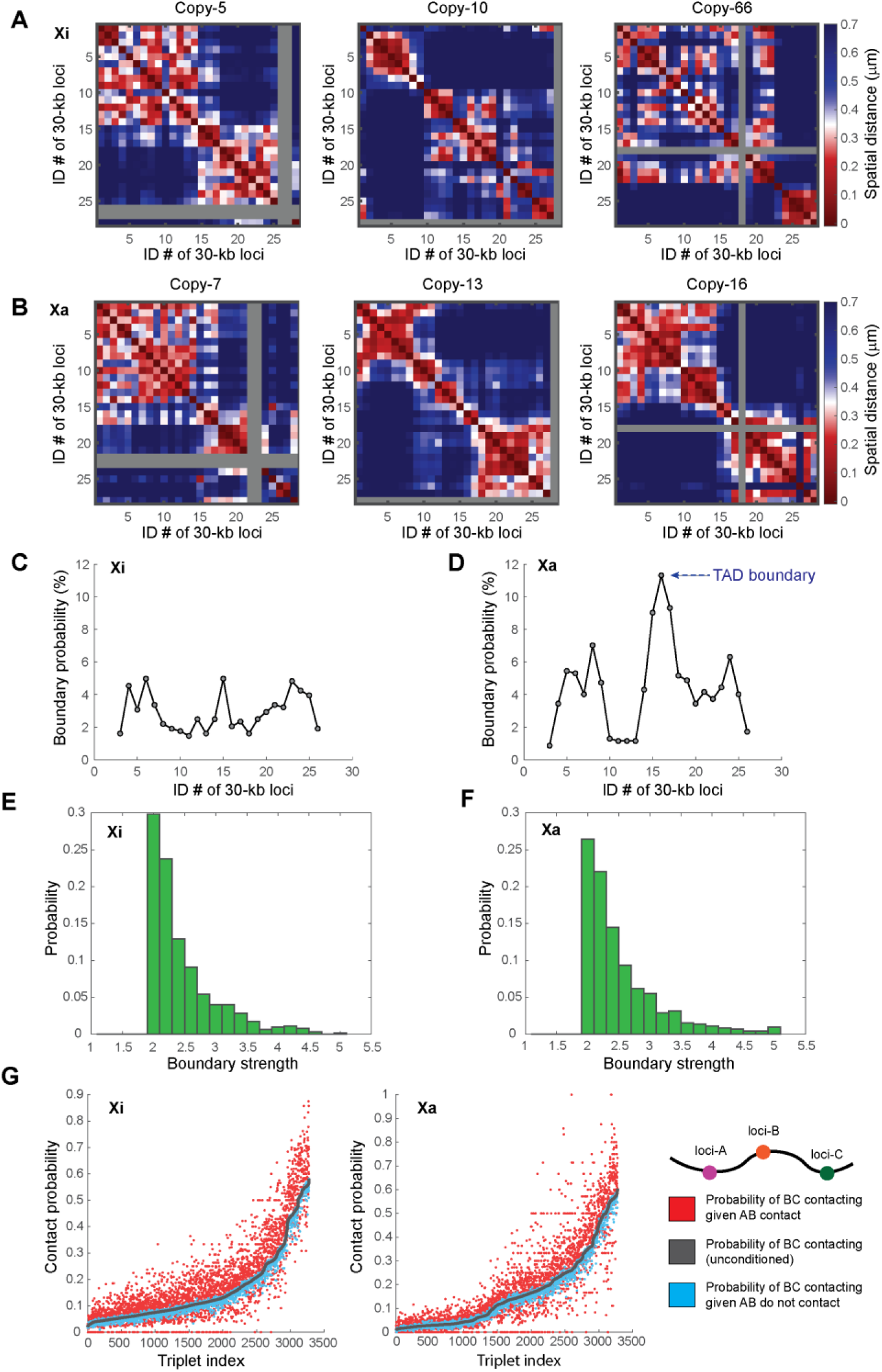
Highly variant TAD-like single-cell domains are present on both inactive and active X chromosomes in IMR-90 cells. (A) Examples of individual spatial distance matrices from single copies of inactive X chromosomes. (B) Examples of individual spatial distance matrices from single copies of active X chromosomes. Gray rows and columns in A and B indicate undetected loci. (C) The boundary probabilities of single-cell domains in inactive X chromosomes. (D) The boundary probabilities of single-cell domains in active X chromosomes. Note the higher boundary probability at the population averaged TAD boundary (arrow). (E) The boundary strengths of single-cell domains in inactive X chromosomes. (F) The boundary strengths of single-cell domains in active X chromosomes. (G) Cooperativity of chromatin interaction in inactive and active X chromosomes. For all ordered triplets in the traced chromatin region, contact probabilities (defined with 200-nm proximity threshold) of genomic loci B and C given contact/non-contact of genomic loci A and B are shown in red and blue, respectively. “Ordered” means that the locus numbers of A, B and C are in an ascending order. The unconditional contact probabilities between loci B and C are shown in gray. The triplet indices are arranged in an ascending order based on the unconditional contact probabilities between loci B and C. N = 342 for C, E and the left panel of G. N = 349 for D, F and the right panel of G.

To check the prevalence of the X chromosome single-cell domains in different cell backgrounds, we imaged another human female cell line, hTERT-RPE-1, derived from retina pigmented epithelium, and found that the above observations in IMR-90 lung fibroblast cells were conserved in the hTERT-RPE-1 cells: Xa showed strong ensemble TAD boundary whereas Xi did not (Fig. 3A-B). Individual Xa and Xi traces showed similarly frequent and strong single-cell domains (Fig. 3C-H). These observations show that the single-cell domains on Xa and Xi are not unique to the IMR-90 cells.

**Figure 3.**
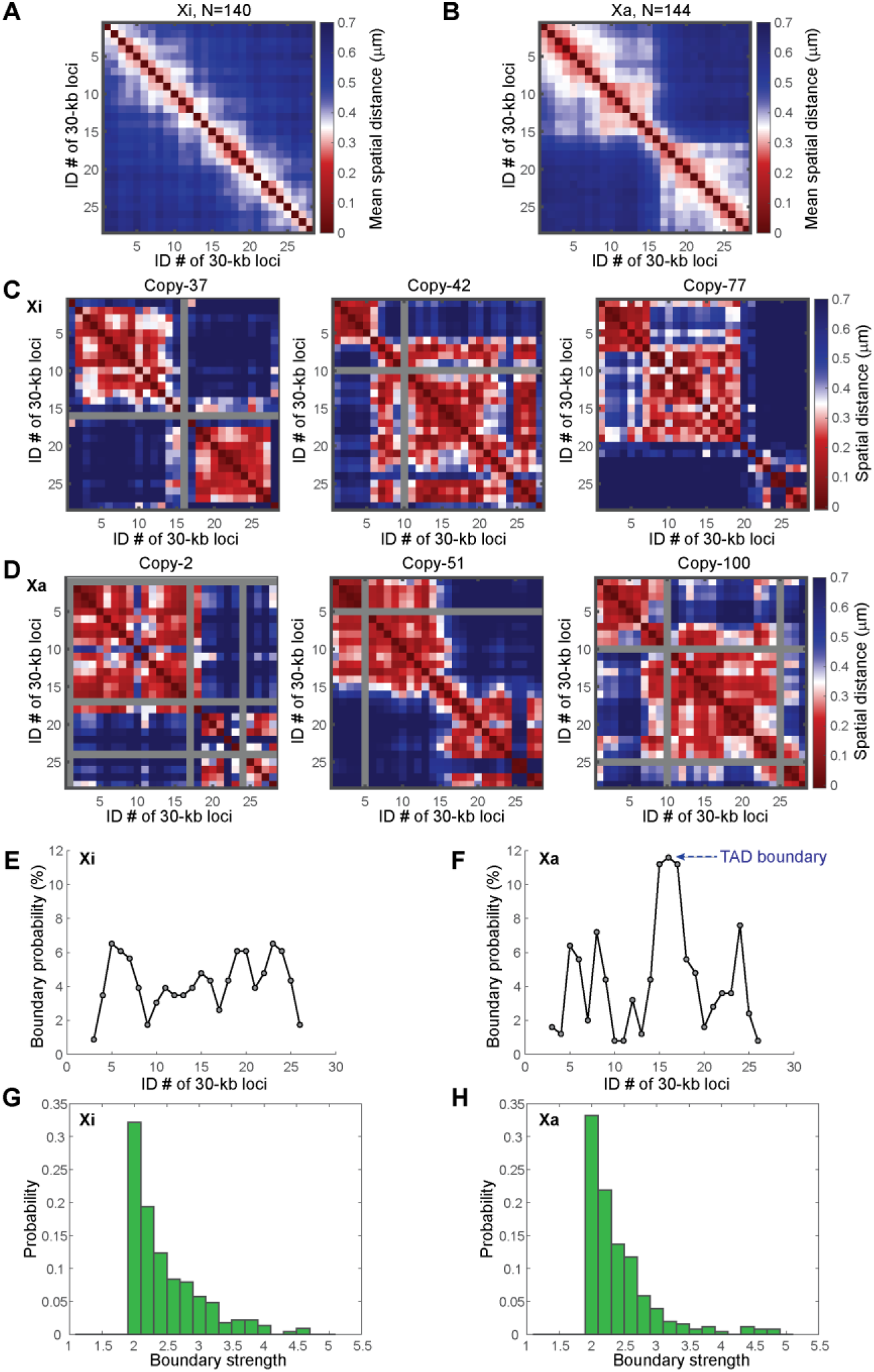
hTERT-RPE-1 cells show similar single-cell domains to those in IMR-90 cells. Data from hTERT-RPE-1 cells are shown in this figure. (A) Mean spatial distance matrix of inactive X chromosomes. (B) Mean spatial distance matrix of active X chromosomes. (C) Individual spatial distance matrices of single copies of inactive X chromosomes. (D) Individual spatial distance matrices of single copies of active X chromosomes. Gray rows and columns in C and D indicate undetected loci. (E) Boundary probabilities in inactive X chromosomes. (F) Boundary probabilities in active X chromosomes. (G) Boundary strengths in inactive X chromosomes. (H) Boundary strengths in active X chromosomes. N = 140 for A, E and G. N = 144 for B, F and H.

Next, to test the hypothesis that epigenetic interactions underlie the formation of single-cell domains, we applied several epigenetic perturbations using commercially available small molecule inhibitors, and measured their effects on the single-cell domains. First we applied histone deacetylase (HDAC) inhibitors trichostatin A (TSA) and sodium butyrate (NaBu) that lead to hyperacetylation of histones, given that histone acetylation prevents nucleosome compaction^20,43,44^. Immunofluorescence staining confirmed the effectiveness of the TSA and NaBu treatment (Fig. S1). Surprisingly, the single-cell domains persisted in both Xa and Xi under the drug treatment with similar boundary frequency and strength as in the untreated control (Fig. 4). We further tested treatments with dimethyloxalylglycine (DMOG) – an HIF-prolylhydroxylase inhibitor, which causes a global increase of histone methylation^45^, GSK126 – an enhancer of zeste homolog 2 (EZH2) methyltransferase inhibitor, and α-amanitin, an RNA polymerase inhibitor. None of the treatments significantly affected the appearance, boundary frequencies, or strengths of the single-cell domains in Xa or Xi (p>0.01 with Bonferroni correction) (Fig. 4B-F, S1). These observations suggest that several major epigenetic components, including histone acetylation and methylation, and RNA polymerase may not be the major factors determining the formation or maintenance of single-cell domains.

**Figure 4.**
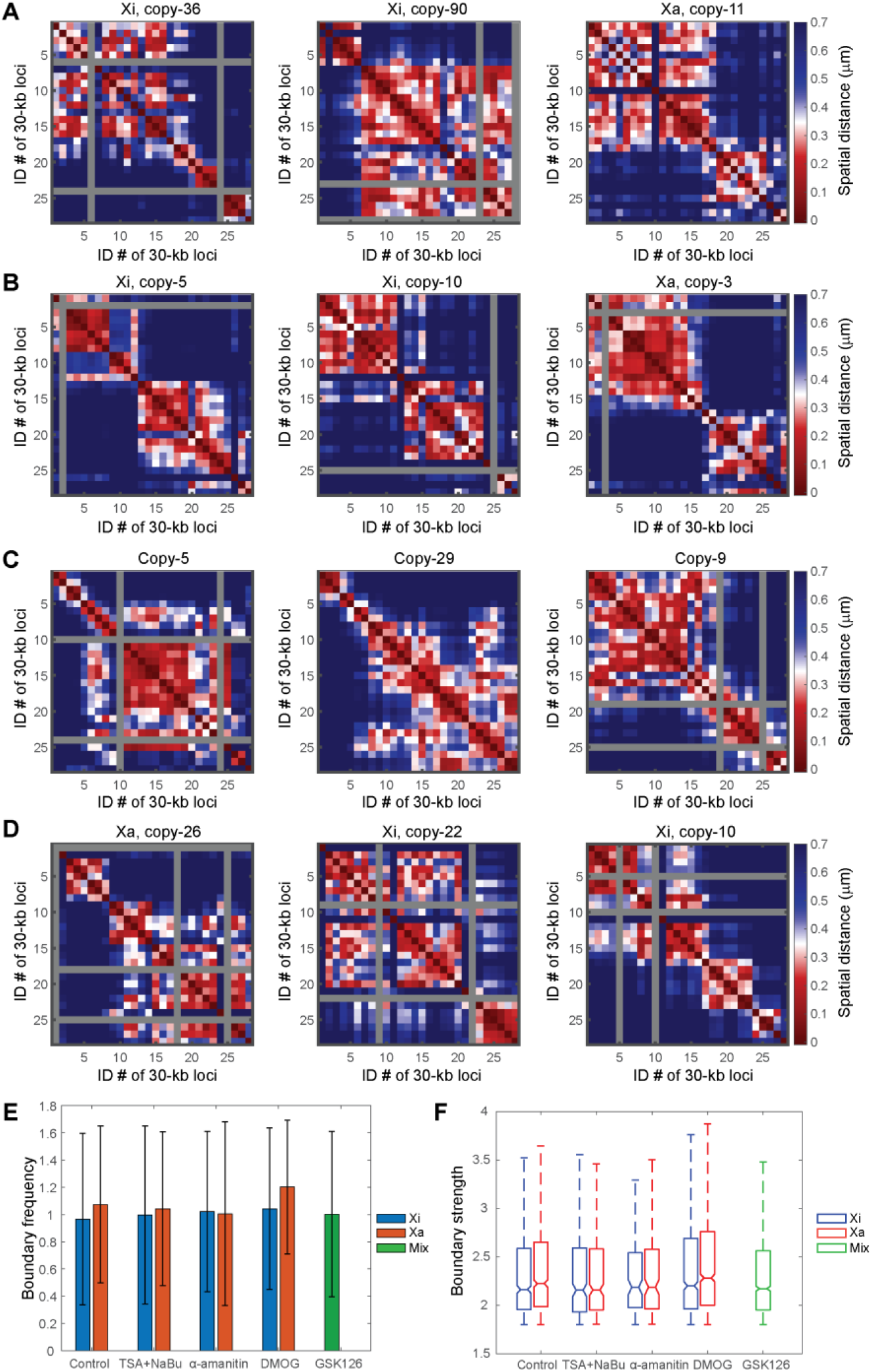
Single-cell domains persist under perturbations of epigenetic components and transcription machinery. (A) Single-cell domains with HDAC inhibitor TSA + NaBu treatment (leading to hyper acetylation of histones). (B) Single-cell domains with DMOG treatment (leading to hyper methylation of histones). (C) Single-cell domains with GSK126 treatment (leading to lower H3K27me3). (D) Single-cell domains with α-amanitin treatment (leading to inhibition of transcription). Gray rows and columns in A-D indicate undetected loci. (E) Average boundary frequencies with or without TSA + NaBu, α-amanitin, DMOG and GSK126 treatments.Error bars stand for standard deviation. (F) Boundary strengths with or without TSA + NaBu, α-amanitin, DMOG and GSK126 treatments. For each box, the horizontal lines from top to bottom represent the non-outlier maximum, 75% quantile, median, 25% quantile and non-outlier minimum. Outliers are defined as values that are more than 1.5 times the interquartile range away from the bottom or top of the box. (In E and F from left to right: N = 342, 349, 132, 154, 68, 84, 347, 473, 884).

Finally, we asked if the increased level of chromatin compaction in Xi in comparison to Xa is present at the scale of single-cell domains. Xi is known to be more spatially compact than Xa at the whole chromosome level^46^. This does not require/imply the compaction to be present/proportional at all length scales. To identify the length scale at which this compaction occurs, we calculated the radii of gyration of the traced genomic regions in Xa and Xi. The comparison showed that the Xa and Xi traces had similar radii of gyration at the traced length scale of single TADs (Fig. 5A). In fact, the Xa traces appear to have slightly smaller radii of gyration than Xi traces in the targeted genomic region (Fig. 5A). However, when larger scale chromosome traces from our previous study^30^ were analyzed in the same fashion, the analyses showed that Xa had significantly larger radii of gyration than Xi at the TAD-to-chromosome length scale (Fig. 5B). These results indicate that the chromatin compaction associated with X inactivation occurs at a length scale above that of TADs and single-cell domains (Fig. 5C).

**Figure 5.**
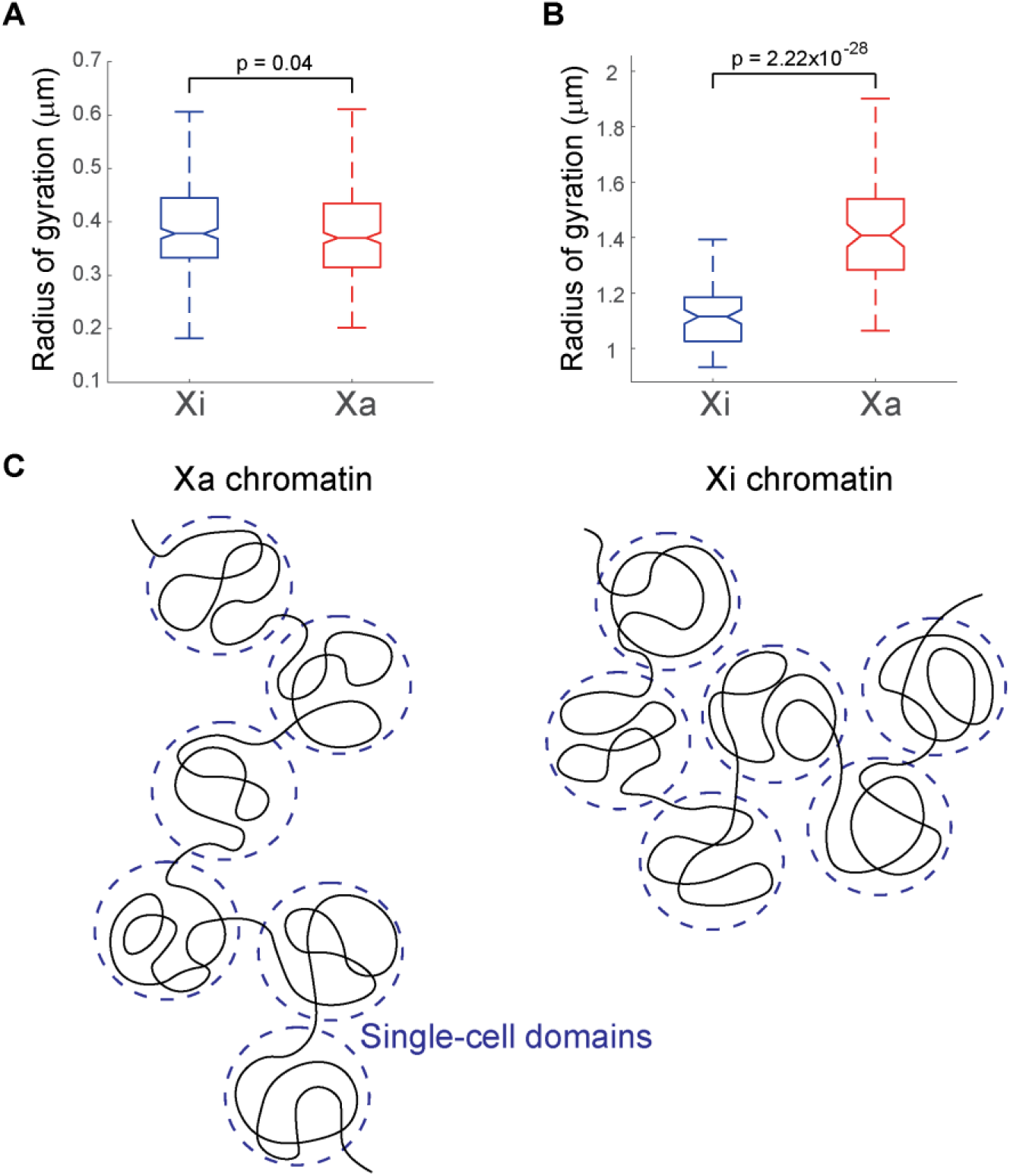
Increased compaction of inactive X chromosomes in comparison to active X chromosomes is present at the TAD-to-chromosome scale, but not at the TAD/sub-TAD scale. (A) Radii of gyration of inactive (left, N=342) versus active X chromosomes (right, N=349) from fine-scale chromatin tracing of the 840-kb region. (B) Radii of gyration of inactive (left, N=95) versus active X chromosomes (right, N=95) from large-scale chromatin tracing of 40 TADs across the entire X chromosome. (C) Schematic illustration of non-proportional compaction of inactive X chromosomes at length scales above the TADs/single-cell domains. The *p* values were calculated for radii of gyration using two-sided Student’s t tests. Boxes are defined as in Fig. 4F.

## Discussion

Loop extrusion and epigenetic landscape have been shown to be major, parallel contributing factors to the spatial organization of TADs^26,27^. As the disruption of loop extrusion did not abolish single-cell domains^32^, recent studies proposed epigenetic interactions mediate the formation of single-cell domains and demonstrated such possibility with computational modeling^34,35^. To recapitulate the highly varied single-cell domains, the modeling involves several constant epigenetic profiles (termed “binding sites” in the model) across a genomic region; different epigenetic reader proteins (termed “binders” in the model) dynamically bind to the corresponding epigenetic binding sites and can mediate the interactions between pairs of genomic loci with the same kind of epigenetic binding sites; the thermodynamics of the chromatin polymer leads to contacts between genomic loci, some of which are captured by the epigenetic interactions mediated by the binders; due to the thermodynamic nature of the system, the distribution of the chromatin contact instances and the epigenetic interaction instances differ from one copy of the polymer to another, which underlies the different single-cell domain structures^34,35^. In this work, we showed that the active and inactive X chromosomes, despite having distinct epigenetic states, both contain single-cell domains of similar features. This observation did not appear to depend on a particular human cell type. A concurrent preprint also reported the observation of single-cell domains in mouse brain tissue, supporting the prevalence of these structures^47^. We further showed that several perturbations of major epigenetic components, including histone acetylation and methylation, and RNA polymerase, did not abolish the single-cell domains. These results did not support the model in which epigenetic interactions mediate single-cell domain formation. In addition, we showed that the chromatin compaction associated with X inactivation occurs at a length scale above that of TADs and single-cell domains.

Due to the limitation of this study, we cannot rule out the possibility that some untested epigenetic components, including various histone and DNA modifications and non-histone DNA binding proteins, underlie the formation of single-cell domains. Neither can we rule out the possibility that the tested epigenetic components are compensated by some untested epigenetic components. Thus this work does not disprove the epigenetic model. However, our data suggest that alternative explanations for the formation of single-cell domains may deserve equal consideration. For example, an unknown motor protein other than cohesin may drive the formation of single-cell domains through loop extrusion. Alternatively, some nuclear components may form scaffolds or aggregates that push away chromatin into adjacent pockets of free nuclear space that correspond to single-cell domains. Further studies may reveal the mechanism underlying the single-cell domains, as well as their dynamics and functional significance in single cells. We also note the recent reports of multiple highly variant chromatin domain structures at the single-cell level observed with different advanced technologies, *e.g.* nucleosome clutches^8^, chromatin nanodomains (CNDs)^20^, packing domains (PDs)^48^, and larger single-cell domains (SCDs) that are multi-megabase in size^49^. The relationship between these structures and the TAD-like single-cell domains identified by chromatin tracing also requires further investigation.

## Conclusions

We conclude that TAD-like structures/single-cell domains exist on both active and inactive X chromosomes and persist under perturbations of major epigenetic components. These observations point towards a mechanism that does not rely on cohesin or major epigenetic marks for the establishment/maintenance of single-cell domains, an apparently prevalent building block of genome architecture.

## Methods

### Probe design and synthesis

To design DNA FISH probes for chromatin tracing, the genomic region of interest (Chr X: 76,800,000 – 77,640,000, hg18) was divided into 28 consecutive 30-kb regions. For each target region, 400 oligonucleotides were designed as template oligos. On each template oligo, the following sequences were concatenated from 5’ to 3’: 1) a 20-nucleotide (nt) forward priming sequence; 2) a 30-nt secondary probe binding sequence; 3) a 30-nt genome targeting sequence; 4) a 20-nt reverse priming sequence. The 30-nt genome targeting sequences were designed with the software OligoArray 2.1^50^ using the following criteria: The melting temperatures of the targeting sequences are between 60 °C and 100 °C; the melting temperatures of potential secondary structures in the sequences do not exceed 70 °C; the melting temperatures of potential cross hybridization among the sequences do not exceed 70 °C; the GC contents of the targeting sequences are between 30% and 90%; there are no consecutive repeats of seven or more A’s, G’s, C’s and T’s; the targeting sequences do not overlap. The chosen targeting sequences were further checked with BLAST+^51^ and only sequences that appear once in the human genome were retained. The secondary probe binding sequences and priming sequences were introduced in previous studies^30,42^. The genomic coordinates of the targeting regions were provided in Supplementary Table 1. The template oligo pool sequences were provided in Supplementary Table 2.

To design *Xist* RNA FISH probes, we downloaded the *Xist* transcript sequences from https://genome.ucsc.edu/cgi-bin/hgGene?hgg_gene=uc004ebm.2&hgg_prot=uc004ebm.2&hgg_chrom=chrX&hgg_start=73040485&hgg_end=73072588&hgg_type=knownGene&db=hg19&hgsid=666371357_9PdT3XyjXSVOpavdVHxoox2IVEBf. We then designed 50 template oligos targeting the transcript. Each template oligo contained (from 5’ to 3’) a 15-nt forward priming sequence, a 30-nt transcript targeting sequence, a 15-nt reverse priming sequence. The 15-nt priming sequences were truncated from previously used priming sequences^30,42^. The 30-nt transcript targeting sequences were designed using the OligoArray 2.1^50^ software with the following parameters: The melting temperatures of the targeting sequences are between 66 °C and 100 °C; the melting temperatures of potential secondary structures in the sequences do not exceed 76 °C; the melting temperatures of potential cross hybridizations among the sequences do not exceed 72 °C; the GC contents of the targeting sequences are between 30% and 90%; there are no consecutive repeats of six or more G’s, C’s, T’s and A’s; the targeting sequences do not overlap. We further checked the sequences of the template oligos with BLAST+^51^ and only retained sequences that appeared once in the human genome. The template oligo pool sequences for *Xist* RNA FISH probes were provided in Supplementary Table 3.

The chromatin tracing template oligo pool was ordered from CustomArray, GenScript. The *Xist* RNA FISH template oligo pool was ordered from Integrated DNA Technologies (IDT), Inc. Primary probes were synthesized from the template oligo pool via limited-cycle PCR, *in vitro* transcription, reverse transcription, alkaline hydrolysis and probe purification^30,52,53^. All PCR primers and dye-labeled reverse transcription primers used in probe synthesis, as well as dye-labeled secondary probes used in the “Secondary probe sequential hybridization” section, were purchased from Integrated DNA Technologies (IDT), Inc. We used Alexa Fluor 647-labeled reverse transcription primers to synthesize chromatin tracing primary probes for datasets collected via “Combined chromatin tracing primary hybridization with mH2A1 immunofluorescence staining” so that all primary probes were labeled with Alexa Fluor 647 fluorophores. We used Cy5-labeled and ATTO 565-labeled reverse transcription primers to synthesize *Xist* RNA FISH probes and chromatin tracing primary probes respectively for datasets collected via “Combined chromatin tracing primary hybridization with *Xist* RNA FISH”. The sequences of the primers and secondary probes were listed in Supplementary Table 4.

### Cell culture

The IMR-90 cell line and hTERT-RPE-1 cell line were purchased from American Type Culture Collection (ATCC, cat. no. CCL-186 for IMR-90 and cat. no. CRL-4000 for hTERT-RPE-1). IMR-90 cells were cultured with Eagle’s MEM (ATCC, cat. no. 30-2003) containing 10% (vol/vol) FBS and 1× penicillin-streptomycin. The hTERT-RPE-1 cells were cultured with DMEM: F12 (ATCC, cat.no. 30-2006) medium containing 10% (vol/vol) FBS and 1× penicillin-streptomycin. For imaging, cells were plated onto 40-mm-diameter, #1.5 coverslips in Falcon 60-mm tissue culture dishes. When the cells reached a confluency of 70%, the cells were fixed with 4% (wt/vol) paraformaldehyde (PFA) in DPBS for 10 min at room temperature, and washed twice with Dulbecco’s phosphate-buffered saline (DPBS).

### Drug treatment

To inhibit transcription, IMR-90 cells were cultured in cell media supplemented with 50 μg/mL α-Amanitin (Sigma-Aldrich, cat.no. A2263) for 8 hours before PFA fixation. To inhibit HDAC function, IMR-90 cells were cultured in cell media supplemented with 100 ng/mL TSA (Sigma-Aldrich, cat.no. T8552) plus 20 mM NaBu (Sigma-Aldrich, cat.no. B5887) for 24 hours before PFA fixation. To inhibitor EZH2 methyltransferase, IMR-90 cells were cultured in cell media supplemented with 1 μM GSK126 (Sigma-Aldrich, cat. no. 5.00580) for 48 hours before PFA fixation. To induce hyper histone methylation, IMR-90 cells were cultured in cell media supplemented with 1 mM or 2.5 mM DMOG (Sigma-Aldrich, cat.no. D3695) for 12 hours before PFA fixation.

### Combined chromatin tracing primary hybridization with mH2A1 immunofluorescence staining

Half of the wild type IMR-90 datasets and all wild type hTERT-RPE-1 datasets were collected with “Combined chromatin tracing primary hybridization with mH2A1 immunofluorescence staining”. The fixed and washed IMR-90 cells or hTERT-RPE-1 cells were treated with freshly made 1-mg/mL sodium borohydride in DPBS for 10 min at room temperature, and washed twice with DPBS for 2 min each. The cells were then permeabilized with 0.5% (vol/vol) Triton X-100 in DPBS for 10 min at room temperature and washed twice with DPBS for 2 min each. Then the cells were treated with 0.1 M HCl for 5 min at room temperature and washed twice with DPBS. Next the cells were treated with freshly prepared 0.1 mg/mL ribonuclease A solution for 45 min at 37 °C. The cells were then washed with 2× saline-sodium citrate (SSC) buffer twice and incubated at room temperature for 30 min in pre-hybridization buffer composed of 50% (vol/vol) formamide and 0.1% (vol/vol) Tween-20 in 2× SSC. Hybridization buffer composed of 50% (vol/vol) formamide and 20% (wt/vol) dextran sulfate in 2× SSC containing 6-20 μM chromatin tracing primary probes was pipetted onto a glass slide. The cell coverslip was flipped onto the glass slide so that cells were in contact with the hybridization buffer. The coverslip-slide assembly was placed onto an 86 °C digital dry bath and denatured for 3 min. The assembly was incubated overnight for 15 – 18 h at 37 °C in a humid chamber. The cells were then washed with 2× SSCT (2× SSC and 0.1% (vol/vol) Tween-20) at 60 °C in a water bath for 15 min twice. The cells were further washed with 2× SSCT at room temperature for 15 min. Next the cells were blocked with 3% (wt/vol) BSA (Sigma-Aldrich, cat. no. A9647-100G) and 0.2% (vol/vol) Triton X-100 in DPBS for 30 min. The cells were then incubated with rabbit mH2A1 primary antibodies (Abcam, ab183041) at a concentration of 1:100 in antibody dilution buffer containing 1% (wt/vol) BSA and 0.2% (vol/vol) Triton X-100 for 1 h at room temperature, and washed three times for 5 min each in DPBS containing 0.05% (vol/vol) Triton X-100. The cells were then incubated with Alexa Fluor 568-labeled goat anti-rabbit secondary antibodies (Invitrogen, A-11011) in antibody dilution buffer for 1 h at room temperature, and washed three times for 5 min each in DPBS containing 0.05% (vol/vol) Triton X-100. Then 0.1-μm yellow-green fiducial beads (Invitrogen, F8803) were resuspended in 2× SSC and applied to the cells so that they could serve as fiducial markers to cancel the sample drift during the sequential hybridization.

### Combined chromatin tracing primary hybridization with *Xist* RNA FISH

Half of the wild type IMR-90 datasets and all drug-treated IMR-90 datasets were collected with “Combined chromatin tracing primary hybridization with *Xist* RNA FISH”. The fixed and washed IMR-90 cells were permeabilized with 0.5% (vol/vol) Triton X-100 in DPBS for 10 min at room temperature and washed twice with DPBS for 2 min each. The cells were then treated with 0.1 M HCl for 5 min at room temperature and washed twice with DPBS. Next, the cells were incubated at room temperature for 30 min in pre-hybridization buffer composed of 50% (vol/vol) formamide and 2 mM Ribonucleoside vanadyl complexes (Sigma-Aldrich, R3380) in 2× SSC buffer. Hybridization buffer composed of 50% (vol/vol) formamide, 0.1% (wt/vol) yeast tRNA (Life Technologies, 15401011), 10% (wt/vol) dextran sulfate (Sigma, D8906-50G) and 100× diluted murine RNase inhibitor in 2× SSC containing 6-20 μM chromatin tracing primary probes and 1 μM *Xist* RNA FISH probes was pipetted onto a glass slide. The coverslip was flipped onto the glass slide so that cells were in contact with the hybridization buffer. The coverslip-slide assembly was placed onto an 86 °C digital dry bath and denatured for 3 min. The assembly was incubated overnight for 15 – 18 h at 37 °C in a humid chamber. The cells were then washed with 2× SSCT at 60 °C in a water bath for 15 min twice, and further washed with 2× SSCT for 15 min and 2× SSC for 3 min at room temperature. The 0.1-μm yellow-green fiducial beads (Invitrogen, F8803) were resuspended in 2× SSC and applied to the cells so that they could serve as fiducial markers to cancel the sample drift during the sequential hybridization.

### Secondary probe sequential hybridization

After the primary probe hybridization, the sample was assembled into a Bioptech’s FCS2 flow chamber and repetitively hybridized with Alexa Fluor 647 and ATTO 565-labeled secondary probes (Supplementary Table 4), imaged and photobleached.

To perform buffer exchange automatically during the secondary probe sequential hybridization procedure, we used a computer-controlled, home-built fluidics system^30,52^. Prior to the secondary probe sequential hybridization, in a first imaging round denoted as hybridization round 0 (hyb 0), for datasets collected with “Combined chromatin tracing primary hybridization with mH2A1 immunofluorescence staining”, we imaged the mH2A1 immunofluorescence patterns and imaged the entire targeted chromatin region with z-stepping in the 560-nm channel and 647-nm channel with the Dual-View setup respectively in 2× SSC. For datasets collected with “Combined chromatin tracing primary hybridization with *Xist* RNA FISH”, we imaged the *Xist* RNA FISH signals and visualized the targeted chromatin region with z-stepping in the 647-nm and 560-nm channels respectively in oxygen scavenging imaging buffer^54^ (50 mM Tris-HCl pH 8.0, 10% (wt/vol) glucose, 2 mM Trolox (Sigma-Aldrich, 238813), 0.5 mg/mL glucose oxidase (Sigma-Aldrich, G2133), 40 μg/mL catalase (Sigma-Aldrich, C30), 0.05% (vol/vol) murine RNase inhibitor in 2× SSC). We covered the oxygen scavenging imaging buffer under a layer of mineral oil (Sigma, 330779) to prevent continuous oxidation. We then photobleached all the signals in hyb 0. For each round of the secondary probe hybridization, the sample was first hybridized with 7.5 nM each of the Alexa Fluor 647 and ATTO 565-labeled secondary probes in 20% (vol/vol) ethylene carbonate (Sigma-Aldrich, E26258) in 2× SSC and incubated at room temperature for 20 min. We then sequentially flowed through the chamber 2 mL wash buffer containing 20% (vol/vol) ethylene carbonate in 2× SSC and 2 mL imaging buffer (either 2× SSC for “Combined chromatin tracing primary hybridization with mH2A1 immunofluorescence staining” experiments or the oxygen scavenging imaging buffer^54^ in “Combined chromatin tracing primary hybridization with *Xist* RNA FISH” experiments).

Next, we took z-stepping images with 647-nm, 560-nm and 488-nm laser illuminations for Alexa Fluor 647 and ATTO 565-labeled secondary probes and fiducial beads respectively, with 200 nm step sizes and 0.4 s exposure time at each step. The z-stacks cover a range of 7 μm in z. After the imaging, we switched the sample buffer to 2× SSC and performed photobleaching with simultaneous 647-nm and 560-nm laser illuminations for 25 s. We performed 14 rounds of secondary hybridization (hyb) in total. The first 14 regions were imaged in the 647-nm channel from hyb 1 to hyb 14 and the last 14 regions were imaged in the 560-nm channel from hyb 1 to hyb 14.

### Image system

For imaging, we used a home-built microscope system with a Nikon Ti2-U body equipped with a Nikon CFI Plan Apo Lambda 60× Oil (numerical aperture: 1.40) objective lens, an active autofocusing system and a Hamamatsu Orca Flash 4.0 V3 camera. The pixel size of our system was 107.9 nm. For epifluorescence illumination, we used a 647-nm laser (2RU-VFL-P-1000-647-B1R, MPB Communications) to excite and image Cy5 and Alexa Fluor 647 fluorophores, a 560-nm laser (2RU-VFL-P-1000-560-B1R, MPB Communications) to excite and image ATTO 565 and Alexa Fluor 568 fluorophores, and a 488-nm laser (2RU-VFL-P-500-488-B1R, MPB Communications) to excite and image the yellow-green fiducial beads for drift correction. For datasets collected with “Combined chromatin tracing primary hybridization with mH2A1 immunofluorescence staining”, we installed a Duel-View setup on the emission path to prevent the fluorescent bleed-through between the mH2A1 and chromatin imaging channels. The Dual-View setup included a 690-nm long-pass dichroic mirror to isolate the emission wavelengths, an FF01-607/70 (Semrock) bandpass filter for fluorescence from the 488-nm and 560-nm channels, and an ET720/60m (Chroma) bandpass filter for fluorescence emissions in the 647-nm channel. We also installed a penta-band notch filter (Chroma, ZET405/488/561/647-656/752m) to block the excitation lasers. To align the 560-nm and 647-nm Duel-View channels and to cancel the color shift, we took z-stack calibration images of 100-nm TetraSpeck beads (Invitrogen). To control the laser intensities of the three lasers above, we used either a filter wheel with neutral density filters (FW102C, Thorlabs) or an acousto-optic tunable filter (AOTF, 97-03309-01 Gooch & Housego) to tune the laser intensities during imaging, and four mechanical shutters (LS3S2Z0, Vincent Associates) to control laser on-off.

### Data analysis

#### Classification of Xa and Xi

For datasets with strong *Xist* FISH signals, we quantified the *Xist* FISH intensity values at the positions of the chromatin trace region in hyb 0 and plotted the distributions of the intensity values. The distribution pattern is typically bimodal due to the enrichment of *Xist* on Xi. We then used the intensity values that separate the two peaks in the distribution as the threshold to classify Xa and Xi. For datasets with mH2A1 immunofluorescence or relatively weak *Xist* FISH signals, to ensure the accuracy of the Xa-versus-Xi classification, we manually selected Xa and Xi traces based on the absence or presence of mH2A1/*Xist* co-localization with the chromatin trace region in hyb 0. We only analyzed cells with two distinct chromatin trace regions in hyb 0 to avoid replicating or non-diploid cells.

#### Reconstruction of 3D chromatin traces

To reconstruct the 3D chromatin traces, we first corrected the color shift between the 560-nm and 647-nm laser channels with TetraSpeck bead images, and measured the sample drift during sequential hybridization and imaging with fiducial bead images. We identified fluorescence loci by local intensity thresholding and measured the 3D positions of each locus with 3D Gaussian fitting in the x, y, z dimensions. The drift-corrected loci were then linked into chromatin traces based on their spatial clustering patterns. If the chromatin traces have missing loci after the initial loci fitting and linking, we attempted to re-fit the 3D positions of the missing loci within the chromatin trace region in corresponding hybridization rounds with 3D Gaussian fitting. The re-fitted loci were then added to the chromatin traces.

#### Hi-C data analysis

We downloaded Hi-C data of IMR-90 cells from Gene Expression Omnibus GSE63525^16^. We first binned the Hi-C contact counts of 5-kb resolution into 30-kb bins, which is the resolution of the chromatin tracing, and summed all contact counts between each pair of traced regions. We defined the summed value as the Hi-C number of reads or contact frequencies.

#### Boundary strength, boundary probability and boundary frequency analyses

Chromatin traces with no less than 25 loci were used for boundary strength, boundary probability and boundary frequency calculations. For each chromatin trace, we reconstructed a spatial distance matrix, with each element representing the Euclidean distance between each pair of traced loci. Missing values were interpolated using linear interpolations of neighboring, non-missing values. To measure the boundary strengths for each locus, we calculated the median values (L) of the left three columns excluding the locus, each extending 10 elements below the diagonal, the median values (R) of the right three 10-element columns including the locus below the diagonal, the median values (T) of the left three 10-element columns including the locus above the diagonal, and the median values (B) of the right three 10-element columns excluding the locus above the diagonal. We then defined two boundary strengths: start-of-domain boundary strength (L/R) and end-of-domain boundary strength (B/T). Using the start-of-domain and end-of-domain boundary strengths, we identified start/end boundary positions by calling the local maximum above a defined threshold. Based on the boundary positions, we computed the start/end boundary probability for each locus as the fraction of chromosomes in which the corresponding locus is identified as a boundary. The average of the start and end boundary probabilities for each locus is defined as the boundary probability of that locus. For each group of chromatin traces, we defined the boundary frequency as the average number of boundaries per trace.

#### Radius of gyration calculation

To calculate the radius of gyration for each chromatin trace, we measured the sum squared Euclidean distance between each traced region to the center-of-mass of the chromatin trace and normalized it to the number of traced regions. The radius of gyration is defined as the square-root of the normalized value. Chromatin traces with no less than 25 detected loci out of 28 targeted loci were used for radius of gyration calculation for fine-scale chromatin tracing. Large-scale chromatin tracing data of 40 TADs across the entire human X chromosome were downloaded from a previous work^30^. Chromatin traces with no less than 35 detected loci were used for radius of gyration calculation for large-scale chromatin tracing.

## Declarations

### Ethics approval and consent to participate

Not applicable.

### Consent for publication

Not applicable.

### Availability of data and materials

The datasets used and/or analyzed during the current study are available from the corresponding author on reasonable request.

### Authors’ contributions

SW conceived study; SW, YC and ML designed the study; YC, ML and MH performed the experiments and analyzed data; SW, YC and ML wrote the paper with help from MH.

## Acknowledgements

We thank Jonathan S. D. Radda for helping with the editing of the manuscript.

---

**Figure S1.**
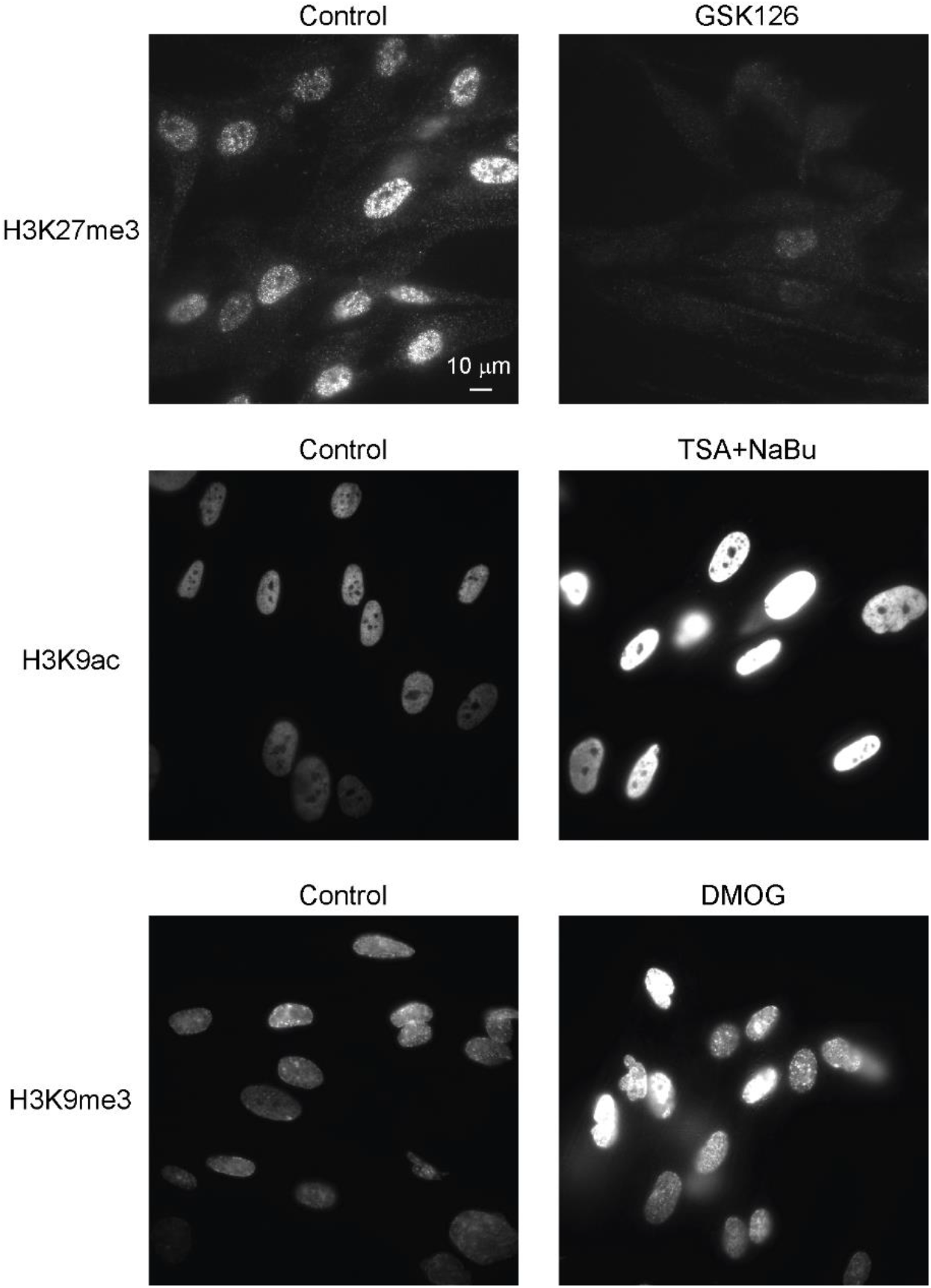
Immunofluorescence staining validation of the effectiveness of drug treatments.

## Supplementary table legends

Supplementary Table 1. The genomic coordinates of the target regions on human X chromosome.

Supplementary Table 2. Sequences of the chromatin tracing template oligo pool.

Supplementary Table 3. Sequences of the *Xist* RNA FISH template oligo pool.

Supplementary Table 4. Sequences of limited-cycle PCR primers, dye-labeled reverse transcription primers and dye-labeled secondary probes.

